# Extracellular matrix chemistry tunes bacterial biofilm metabolism and optimizes fitness

**DOI:** 10.1101/2025.09.18.677115

**Authors:** Jinyang Li, Georgia R. Squyres, Kathy Duong, Courtney Reichhardt, Matthew R. Parsek, Dianne K. Newman

**Affiliations:** Division of Biology and Biological Engineering, Caltech, Pasadena, CA 91125, USA; Division of Geological and Planetary Sciences, Caltech, Pasadena, CA 91125, USA; Department of Chemistry, Washington University, St. Louis, MO 63130, USA; Department of Microbiology, University of Washington, Seattle, WA 98195, USA

**Keywords:** actin dynamics, treadmilling, phase separation, cofilin, VASP

## Abstract

Chemically complex extracellular matrices define cellular microenvironments and shape cell behavior. We hypothesized a composition–properties–function relationship in these natural living materials, where interactions among matrix components govern material properties and cellular physiology. Using *Pseudomonas aeruginosa* biofilms as a model system, we show that electrostatic interactions between the cationic polysaccharide Pel and extracellular DNA (eDNA) regulate retention of pyocyanin (PYO), a redox-active metabolite that supports anaerobic metabolism via extracellular electron transfer (EET). Biofilm-mimetic hydrogels and natural biofilms revealed that altering Pel’s charge via pH adjustment or chemical acetylation, or tuning the Pel:eDNA ratio, predictably modulates PYO retention and EET efficiency. Functionally, a lower Pel:eDNA ratio enhances metabolism under oxygen limitation, whereas a higher ratio promotes survival under antibiotic stress. These findings highlight how matrix chemistry encodes tunable material properties that confer biofilm fitness advantages and establish a materials-based framework for understanding extracellular matrices in multicellular communities.

## Introduction

In nature, cells are almost invariably embedded in self-produced extracellular matrices, creating natural living materials that form the structural and functional foundation of most biological systems.^1–3^ Like engineered materials, these evolutionarily-derived ensembles exhibit intricate compositions and diverse physicochemical properties tailored for specialized functions. Insights into these biological materials have long inspired developments in materials engineering (e.g., bioinspired, biomimetic, and engineered living materials),^4–8^ yet materials science is only beginning to advance our understanding of biology. ^9–11^ Drawing inspiration from the well-established composition–properties–function relationships in classical materials science and engineering, we sought to understand whether analogous principles could be applied to decipher how chemical composition governs material properties and, ultimately, biological functions in natural living materials.

We chose to examine biofilms—materials assembled by nature’s most accomplished molecular engineers: bacteria. Biofilms, where multicellular bacterial assemblies are embedded within a self-produced exopolymeric matrix, dominate microbial life in both environmental and clinical settings.^12^ This matrix provides the structural and functional framework for a remarkably flexible and adaptive mode of life, exhibiting emergent properties not seen in free-swimming cells (e.g., tolerance against various stressors).^13–15^ As such, this matrix is central to the resilience and evolutionary success of biofilms, and understanding it has far-ranging applications, such as being able to remove biofilms from medical implants and overcome their antibiotic tolerance. While considerable progress has been made in identifying and characterizing individual matrix polymers, a critical and often underappreciated perspective is to view the biofilm as an integrated materials system where polymers interact in complex and dynamic ways to define the matrix’s physicochemical properties, and, in turn, shape microbial behavior and physiology.^16,17^

Towards this end, we chose *Pseudomonas aeruginosa* (*P. aeruginosa*)—an opportunistic bacterial pathogen—as our model organism. It is known for forming antibiotic-tolerant biofilms that contribute to both acute and chronic infections.^18,19^ One of the most extensively studied laboratory strains, PA14, forms biofilms with a matrix primarily composed of Pel, an aminopolysaccharide, and extracellular DNA (eDNA).^20^ Prior work has shown that Pel binds eDNA through electrostatic attraction;^21,22^ likewise, eDNA binds to pyocyanin (PYO), a redox-active phenazine metabolite that supports biofilm metabolism when O^2^ is scarce.^23^ This network of interactions, combined with the ease of genetically manipulating *P. aeruginosa*, offers a tractable system to examine how polymer interactions and their relative abundance influence material properties and biofilm metabolism.

The specific material property we examine in this study is facilitation of extracellular electron transfer (EET), a widespread survival strategy biofilm cells use to overcome oxygen limitation.^24^ Cells living in the interior of biofilms experience little to no oxygen, due to the consumption of oxygen by cells on the biofilm periphery outpacing its diffusion to the interior.^25^ To maintain electron flow and sustain energy conservation when oxygen is scarce, cells can employ diffusible redox-active molecules (e.g. PYO), to shuttle electrons derived from intracellular metabolism through the biofilm to oxygen at a distance, a process termed EET. This process maintains anaerobic metabolism of cells in the biofilm core and contributes to the overall resilience of biofilms against environmental stressors.^26,27^

In this study, we investigated how the polymeric composition of the biofilm matrix regulates its EET properties, thereby shaping biofilm metabolism and fitness. Specifically, we asked whether Pel competes with PYO for eDNA binding, and how Pel-eDNA interaction influences PYO retention and biofilm EET. We further examined whether varying the Pel:eDNA ratio modulates EET, and how these changes affect metabolic activity under different environmental conditions. We hypothesized that matrix chemistry can be fine-tuned to adjust material properties and optimize biological performance, a potentially generalizable biofilm survival strategy. More broadly, our findings reveal how the application of concepts from materials science and engineering can illuminate, and even help predict, matrix-driven biological behaviors.

## Results and discussion

### Pel-DNA binding competes with DNA intercalation

To test the hypothesis that Pel competes with the intercalation between PYO and DNA, we first isolated secreted Pel from the supernatant of a genetically engineered Pel-overproducing *P. aeruginosa* strain and confirmed its chemical structure by solid-state ^13^C NMR, finding it to be consistent with previously published data (**Figure 1A**).^21,22,28^ We then examined Pel–DNA interactions by incubating Pel with DNA at pH 5 and pH 8. A visible precipitate was observed only at pH 5 (**Figure S1**), indicating Pel–DNA complex formation under acidic conditions. This result is consistent with previous reports ^21,22^, as well as with our expectation that Pel binds DNA efficiently only below its pKa (previously estimated as ∼6.8)^22^, where it is predominantly protonated and cationic.

**Figure 1.**
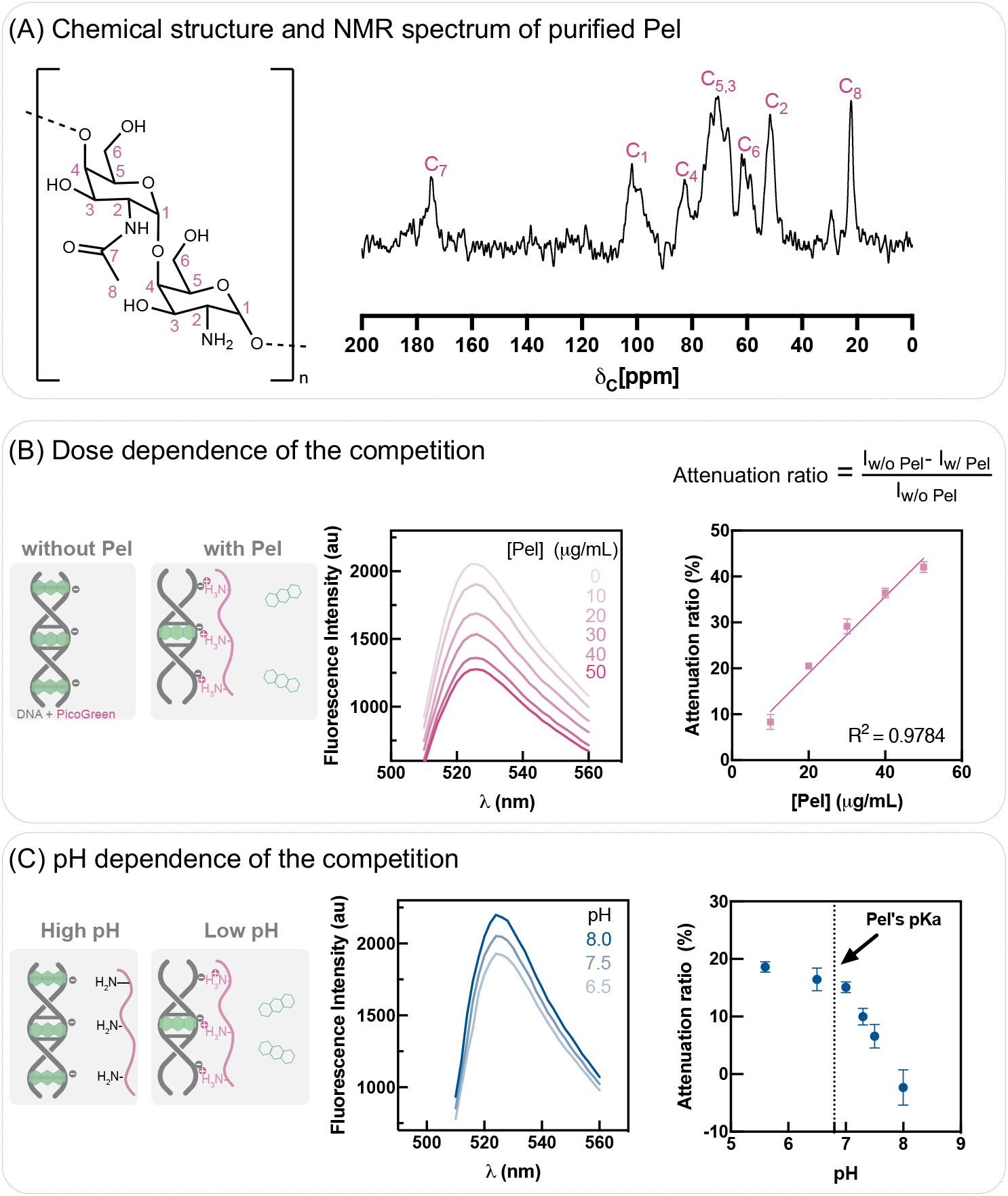
Pel competes with DNA intercalation. **(A)** Pel (structure, left) was purified from a Pel-overproducing strain and verified by ^13^C NMR (right). **(B)** Pel addition inhibits PicoGreen–DNA binding in a dose-dependent manner. Left: schematic shows Pel competing with PicoGreen for DNA binding, reducing fluorescence. Middle: fluorescence spectra show decreased intensity with increasing Pel. Right: attenuation ratios plotted against Pel concentration exhibit a linear relationship. **(C)** The inhibition is enhanced at lower pH. Left: schematic shows increased Pel–DNA binding at low pH, reducing fluorescence from PicoGreen-DNA binding. Middle: representative spectra show greater attenuation at low pH. Right: attenuation ratios increased at lower pH. Attenuation ratio was calculated as the fluorescence difference at 526 nm with and without Pel, normalized to no-Pel intensity at the same pH. The points and error bars indicate mean and standard deviation from triplicates, respectively.

Attempts to directly quantify the competitive binding of Pel and PYO to DNA using isothermal titration calorimetry (ITC) were hindered by Pel’s poor solubility. Instead, we used PicoGreen as a surrogate for PYO, as it also intercalates into DNA but fluoresces only when bound. We hypothesized that if Pel competes with intercalators for DNA binding, introducing Pel would reduce PicoGreen fluorescence. Consistent with this prediction, adding Pel to DNA–PicoGreen mixtures at pH 6.5 caused a dose-dependent, linearly correlated decrease in fluorescence (**Figure 1B**), indicating that Pel binding displaces PicoGreen from DNA.

We further predicted that competition would intensify at lower pH due to stronger Pel–DNA electrostatic interactions. Indeed, fluorescence attenuation increased as pH decreased (**Figure 1C**), consistent with enhanced Pel protonation, stronger Pel–DNA binding, and reduced PicoGreen–DNA binding. Together, these results demonstrate that Pel competitively displaces PicoGreen from DNA. Given that PicoGreen

**Figure S1.**
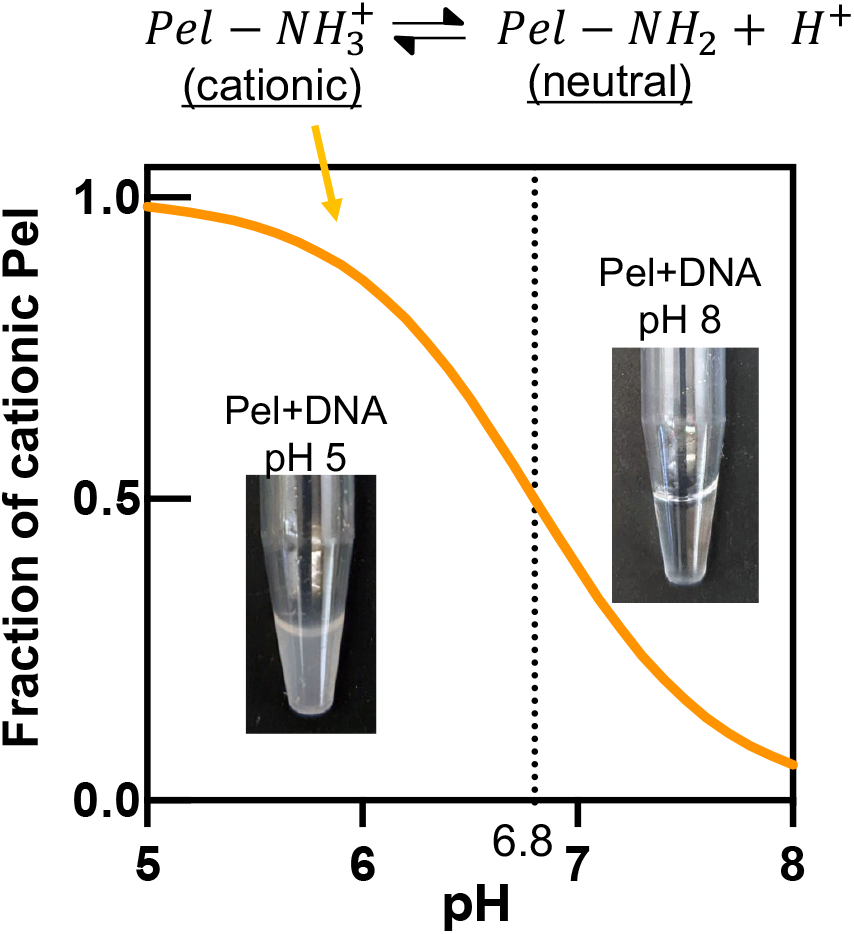
Pel binds to DNA in a pH-dependent way. The orange curve represents the fraction of cationic Pel (pKa = 6.8) predicted by the Henderson-Hasselbalch equation. Insets illustrate this effect: at pH 5, Pel–DNA binding produced a cloudy solution, whereas at pH 8, the absence of binding yielded a clear solution.

### Pel-DNA binding affects PYO retention and EET

That Pel competes with intercalators to bind DNA led us to hypothesize that tuning the interaction between Pel and eDNA would regulate PYO retention and extracellular electron transfer (EET). We first sought to enhance Pel–DNA binding by lowering the pH, reasoning that stronger Pel–DNA interactions would reduce PYO-DNA binding, thereby decreasing PYO retention and diminishing EET efficiency (**Figure 2A**). To test this prediction, we employed an electrochemical approach using an interdigitated electrode array (IDA). The IDA features two closely spaced working electrodes that enable electron transfer between them. Although traditionally used to study the conductivity of abiotic materials, it has been extended to study EET through microbial biofilms^30–33^, and our laboratory recently adapted this system to assess biofilm EET efficiency quantitatively.^23^ Two key parameters are measured: D_loss_, the diffusion coefficient of PYO out of the biofilm, which reflects PYO retention; and D_ap_/D_loss_, which indicates EET efficiency by comparing the apparent electron diffusion coefficient to the molecular diffusion coefficient of PYO (e.g., no EET when D_ap_/D_loss_ = 1).

**Figure 2.**
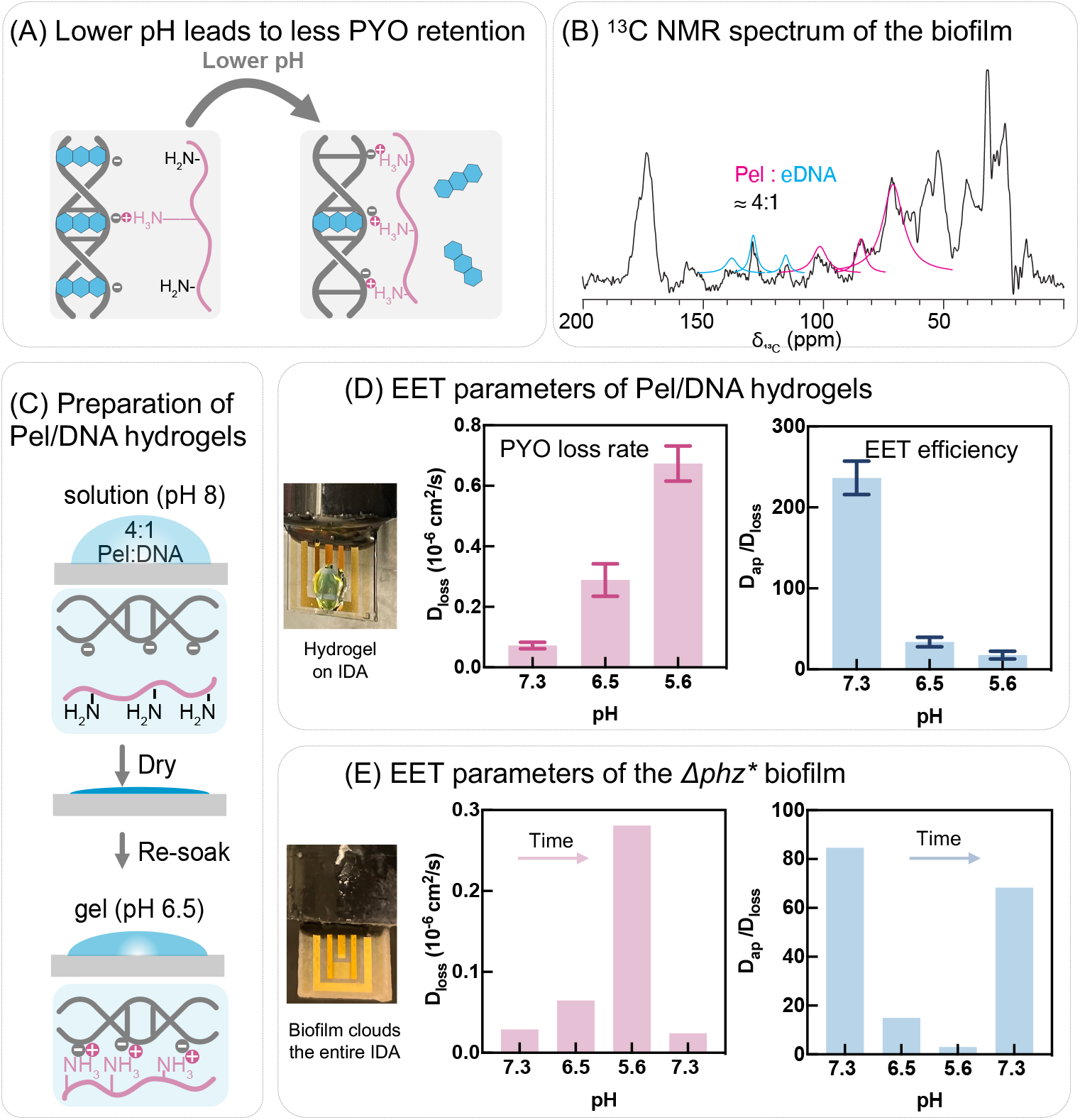
pH tunes PYO retention and EET. **(A)** Schematic: low pH strengthens Pel–DNA binding, reducing PYO–DNA binding, leading to decreased PYO retention and lower EET efficiency. **(B)** ^13^C NMR analysis of biofilm polymeric composition reveals a Pel:eDNA ratio of 4:1. **(C)** Preparation of biofilm-mimetic hydrogels composed of Pel and DNA. **(D)** Pel/DNA hydrogels (4:1) show faster PYO loss and less efficient EET at low pH. Bars represent mean ± SD (n = 3). The leftmost photograph shows representative hydrogels on an IDA electrode (blue due to PYO soaking). **(E)** *Δphz\** biofilms also exhibit faster PYO loss and less efficient EET at low pH, which is reversible upon pH increase; the leftmost photograph shows biofilms grown on an IDA electrode (for higher resolution photos of biofilms on IDAs, see Figure 4D). Data are from a single biofilm, representative of biological triplicates, with triplicate datasets shown in Figure S2.

To test our hypothesis in a well-defined yet physicochemically relevant system, we fabricated a biofilm-mimetic hydrogel. We first used ^13^C NMR to quantify the ratio between Pel and eDNA within PA14 biofilms, the two major matrix components^20^. As described in Methods, the biofilm spectrum, after subtraction of the spectrum obtained from a planktonic culture of the *Δpel* mutant (which does not produce Pel), revealed a Pel:eDNA ratio of approximately 4:1 within the biofilm matrix. (**Figure 2B**). Guided by this finding, we deposited hydrogels composed of Pel and DNA at a 4:1 ratio. onto IDA electrodes. **Figure 2C** illustrates that gel formation was driven by pH-dependent Pel–DNA crosslinking: a solution containing the Pel–DNA mixture at pH 8 was deposited on an IDA and lowering the pH to 6.5 triggered hydrogel formation. The gel-coated electrodes were then soaked in PYO at a defined pH, transferred to PYO-free medium of the same pH, and PYO equilibration into the fresh medium was monitored electrochemically over time, as previously described.^23^ This workflow allowed measurement of EET parameters under different pH conditions. As predicted, Pel/DNA hydrogels exhibited higher D_loss_ and lower D_ap_/D_loss_ at lower pH, indicating reduced PYO retention and less efficient EET (**Figure 2D**).

To validate these results in natural biofilms, we grew PA14 *Δphz\** biofilms on IDA electrodes and performed EET measurements as previously described.^23^ The *Δphz\** mutant, which is incapable of phenazine biosynthesis, was used to eliminate interference from other phenazine species. Biofilms were grown by incubating IDAs in reactors under stirred, oxic conditions. Because intrinsic heterogeneity can lead to substantial biofilm-to-biofilm variability, we focused on comparing EET efficiency within the same biofilms under different pH conditions, thereby controlling for inter-biofilm variation. As shown in **Figure 2E** (with triplicate data shown in **Figure S2**), lowering the pH increased D_loss_ and decreased D_ap_/D_loss,_ consistent with the biomimetic hydrogel results. Furthermore, both parameters nearly returned to their original values when the pH was restored to 7.3, demonstrating that the pH effect on EET is reversible and supporting the conclusion that it is driven by pH-dependent Pel–DNA binding. Notably, the pH conditions tested are within the physiological range, as previous work has shown that the core of *P. aeruginosa* biofilms can reach pH values as low as 3.5.^34,35^

**Figure S2.**
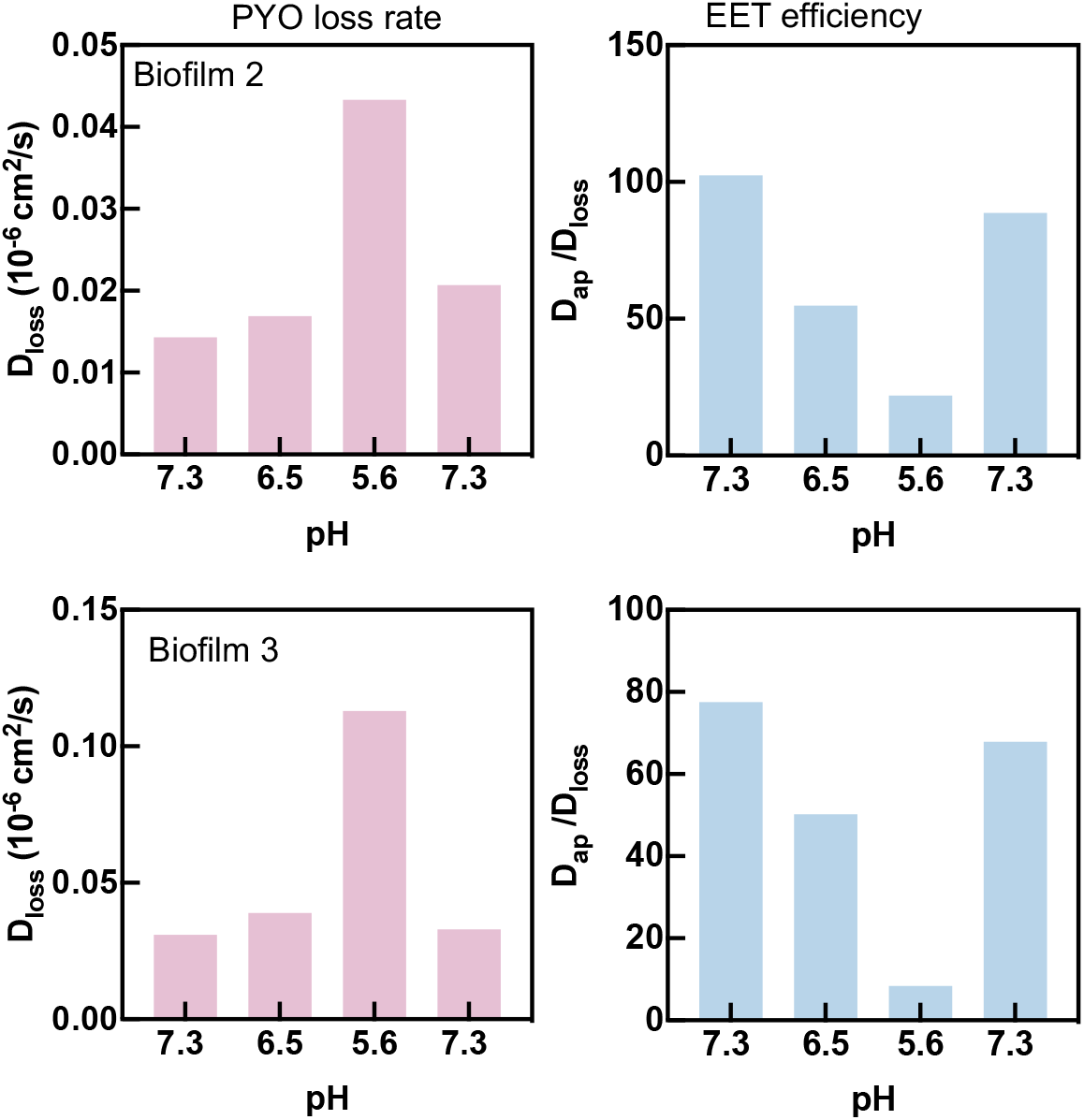
Triplicate datasets for Figure 2E, the pH dependence of biofilm EET.

Next, we employed an orthogonal chemical strategy to perturb Pel–DNA interactions. We hypothesized that acetylation of Pel’s amine groups would neutralize its cationic charges, thereby weakening Pel–DNA binding, increasing PYO–DNA association, enhancing PYO retention, and ultimately improving EET efficiency. To test this, we acetylated both Pel/DNA hydrogels and Δphz* biofilms with acetic anhydride, as previously described,^28,36^ and compared their EET parameters before and after acetylation at pH 6.5 (**Figure 3A)**. As predicted, Pel/DNA hydrogels showed a lower D_loss_ and higher D_ap_/D_loss_ after acetylation, indicating enhanced PYO retention and more efficient EET (**Figure 3B**). A similar trend was observed in *Δphz\** biofilms (**Figure 3C**).

**Figure 3.**
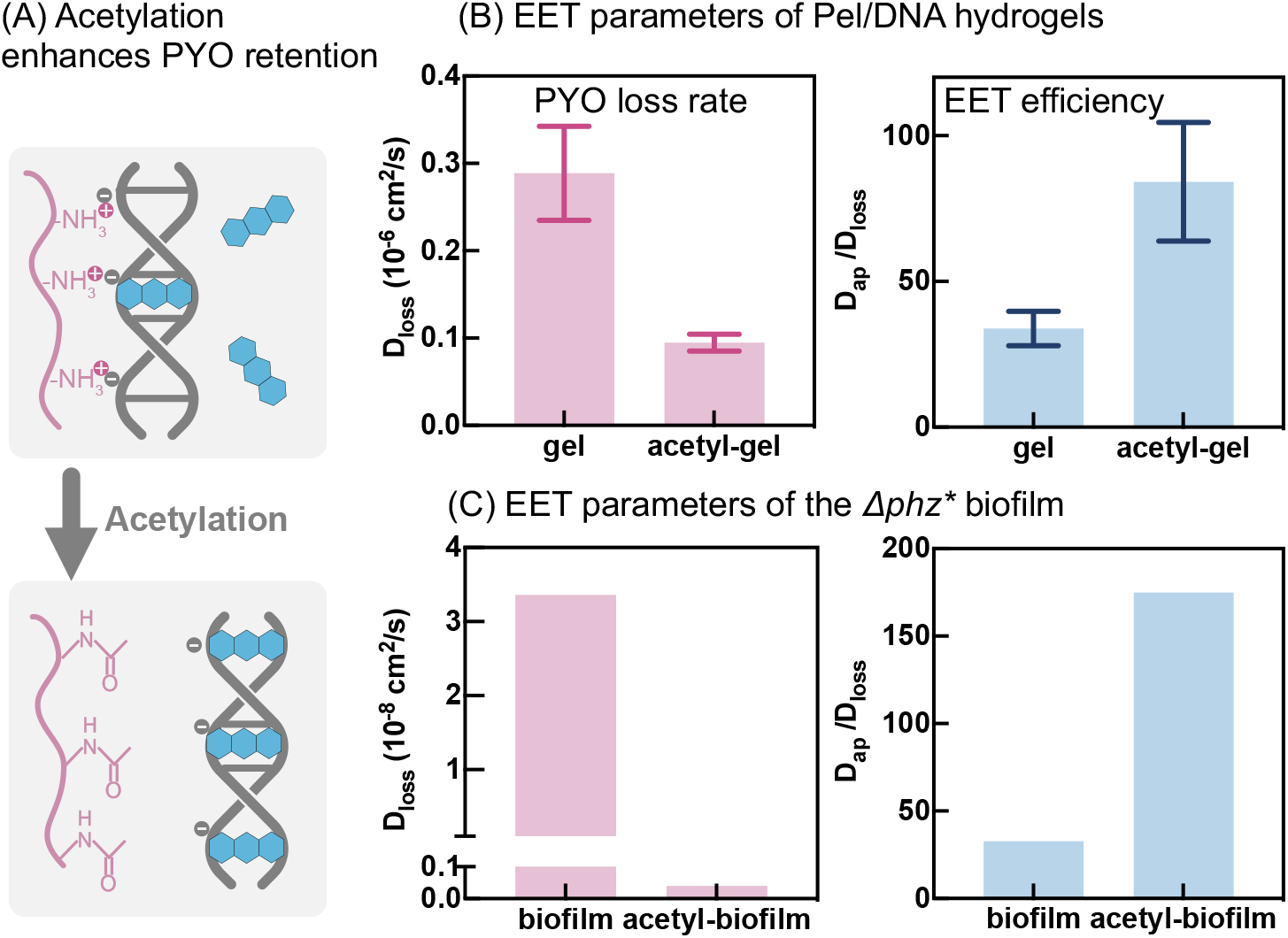
Acetylation enhances PYO retention and EET. **(A)** Schematic: acetylation neutralizes Pel’s amine groups, reducing Pel–DNA binding and allowing increased PYO–DNA binding, which enhances PYO retention and EET. **(B)** Pel/DNA hydrogels show slower PYO loss and higher EET efficiency after acetylation. Bars represent mean ± SD (n = 3). **(C)** *Δphz\** biofilms also exhibit slower PYO loss and enhanced EET after acetylation; data are from a single biofilm, before and after acetylation, representative of biological triplicates; triplicate datasets in Figure S3.

To test whether the acetic anhydride reaction indeed acetylated Pel’s amine groups, we used fluorescently labeled Wisteria floribunda lectin (WFL)^22,37^, which specifically binds N-acetylgalactosamine (GalNAc) but not galactosamine (GalN)^38^. Because Pel is predominantly composed of repeating GalN– GalNAc dimers, acetylation is expected to convert GalN to GalNAc, thereby enhancing WFL binding. We also used TOTO-1 to fluorescently label eDNA.^39,40^ As shown in **Figure S4**, acetylated biofilms exhibited markedly higher WFL fluorescence than untreated biofilms, but comparable TOTO-1 fluorescence. These results indicate that the acetylation reaction modifies Pel without affecting eDNA.

Taken together with the results shown in **Figure 2**, these findings demonstrate that modulating Pel– DNA interactions—enhancing by lowering pH or weakening by acetylation—predictably alters PYO retention and EET. Furthermore, the use of biofilm-mimetic Pel/DNA hydrogels, which isolate biopolymers from cellular factors, indicates that the interactions between Pel and DNA are necessary and sufficient to explain this effect.

**Figure S3.**
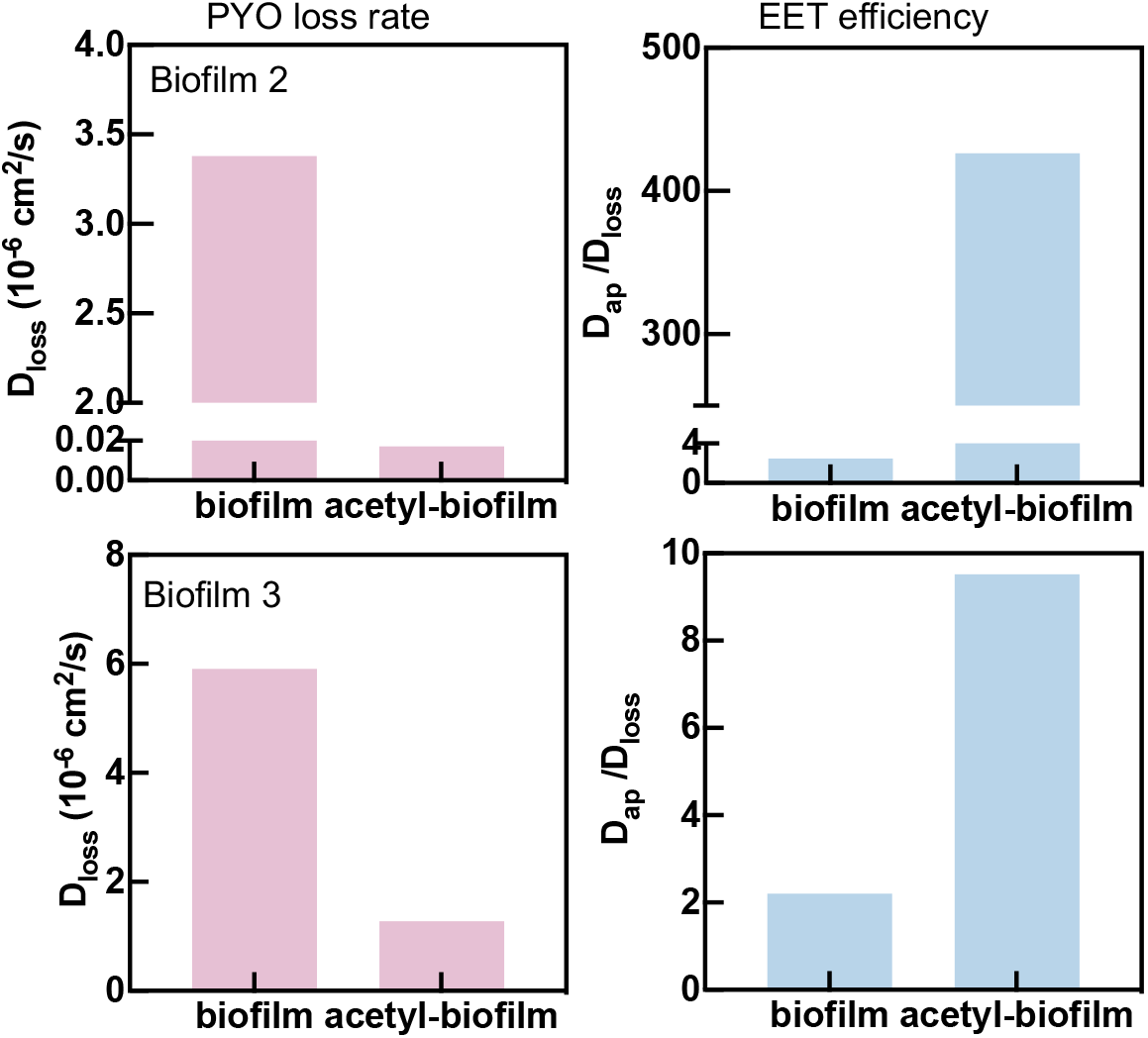
Triplicate datasets for Figure 3C, showing that acetylation enhances biofilm EET.

**Figure S4.**
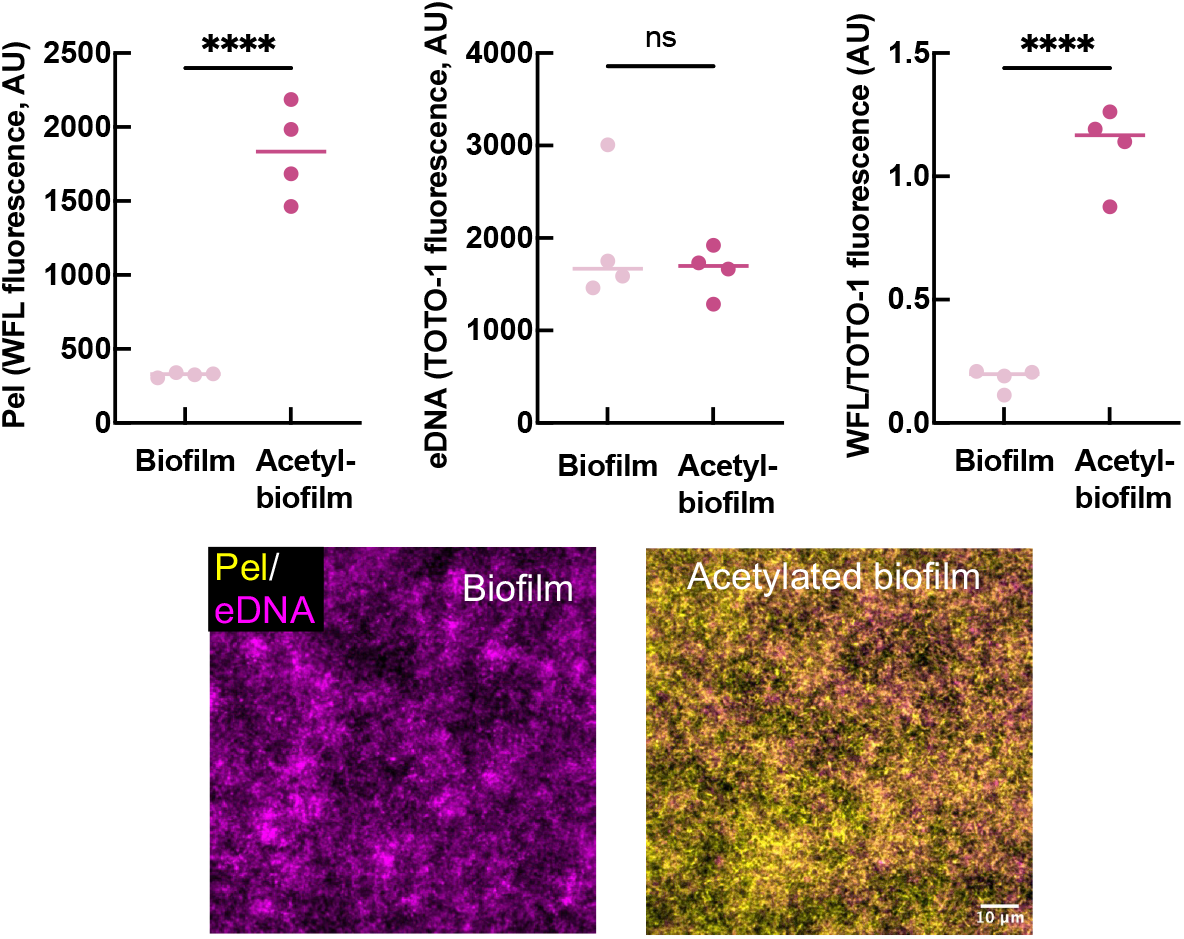
The acetylation of biofilm acetylates Pel’s GalN moieties. TOTO-1 was used to stain eDNA, and WFL, which binds Pel with increased affinity after acetylation, was used to assess Pel acetylation. Top: Fluorescence intensities of WFL, TOTO-1, and their ratio in untreated and acetylated biofilms. P values were calculated using unpaired, two-tailed t test (ns>0.05, ****p<0.0001). Bottom: Representative fluorescence images showing average-intensity projections of confocal Z-stacks.

### Pel:DNA ratio affects PYO retention and EET

Next, we investigated whether the ratio of Pel to eDNA regulates PYO retention and EET in natural biofilms. We predicted that increasing Pel content would lead to more Pel–DNA binding, thereby reducing PYO–DNA binding, resulting in lower PYO retention and diminished EET efficiency (**Figure 4A**). Prior to manipulating natural biofilms, we fabricated biofilm-mimetic hydrogels containing a fixed amount of DNA but varying amounts of Pel. As expected, gels of higher Pel:DNA ratio exhibited higher D_loss_ and lower D_ap_/D_loss_ at pH 6.5, indicating reduced PYO retention and less efficient EET (**Figure 4B**).

**Figure 4.**
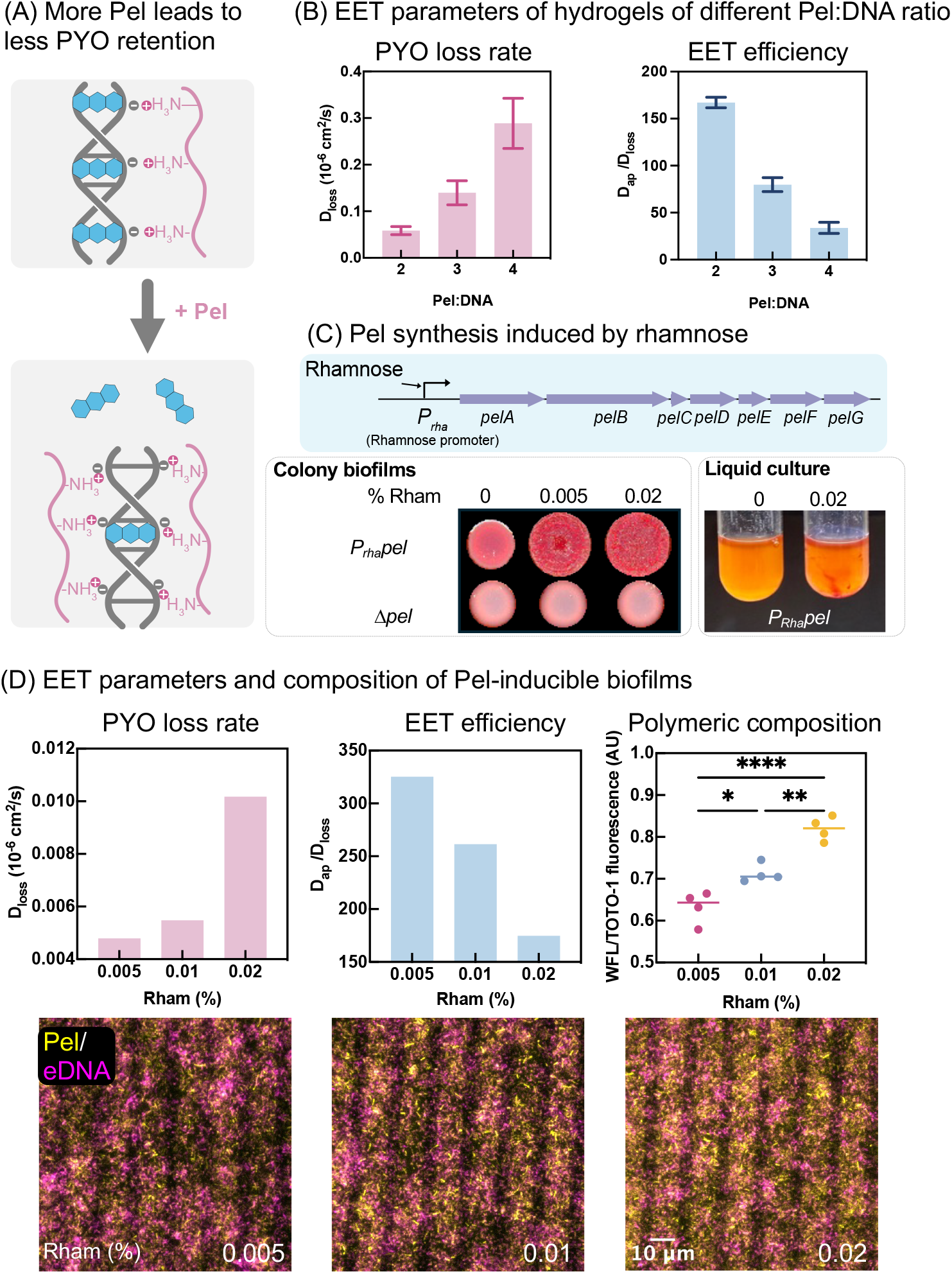
Pel:eDNA ratio regulates PYO retention and EET. **(A)** Schematic: increasing Pel enhances Pel– DNA binding, reducing PYO–DNA binding, which decreases PYO retention and EET efficiency. **(B)** Pel/DNA hydrogels with higher Pel:DNA ratios exhibit faster PYO loss and less efficient EET. Bars represent mean ± SD (n = 3). **(C)** Rhamnose induction up-regulates Pel production in the *P*_*rha*_*pel* strain. Rhamnose-induced *P*_*rha*_*pel* reveals a red, wrinkled colony morphology in colony biofilm (left), and visible red aggregates in the liquid culture (right). Both experiments were stained with Congo Red, a Pel-binding dye, for visualization **(D)** Top: Biofilms induced with more rhamnose display faster PYO loss, reduced EET efficiency (left and middle), and increased Pel:eDNA ratio (right). Data are representative of biological triplicates, with the triplicate datasets shown in Figure S5. P values were determined by one-way ANOVA (ns>0.05, *≤0.05, **≤0.01, ***≤0.001, ****≤0.0001). Bottom: Representative fluorescence images show *P*_*rha*_*pel* biofilms stained with WFL (Pel) and TOTO-1 (eDNA), shown as average-intensity projections of confocal Z-stacks.

To examine this relationship in biofilms in a controlled manner, we engineered a inducible Pel-producing *P. aeruginosa* strain in which expression of the Pel biosynthesis operon *pelA-G* is driven by a rhamnose-inducible promoter. We constructed this strain in the Δ*phz\** background and hereafter refer to it as *P*_*rha*_*pel*. By adjusting the rhamnose concentration, we could modulate Pel production in biofilms (schematic in **Figure 4C**). We also constructed a Pel-nonproducing Δ*pelA-G* knockout in the Δ*phz\** background, which we refer to as Δ*pel*. Two simple tests of Pel overproduction confirmed that our construct was working as expected: (i) when grown on solid agar supplemented with Congo Red, a dye known to bind Pel,^20^ the *P*_*rha*_*pel* strain exhibited a wrinkled and Congo Red-binding colony morphology under rhamnose induction, a phenotype associated with Pel overproduction, whereas the *Δpel* strain formed smooth and pale colonies^28^ (left image, **Figure 4C**); (ii) when cultured in liquid medium containing Congo Red, visible red precipitate was observed for the *P*_*rha*_*pel* strain under rhamnose induction, reflecting crosslinking between Pel and eDNA^22^ (right image, **Figure 4C**). Together, these results establish that rhamnose effectively modulates the production of Pel in the *P*_*rha*_*pel* strain.

To examine EET parameters, biofilms were grown on IDA electrodes in an anaerobic chamber with KNO_3_ as the terminal electron acceptor. This condition was chosen because it produced biofilms of similar thickness across different rhamnose concentrations, whereas growth in oxic bioreactors resulted in *P*_*rha*_*pel* biofilms with substantially different thicknesses which would have confounded the interpretation of the experiments. EET was measured at pH 6.5, and, as predicted, biofilms induced with higher rhamnose concentrations showed higher D_loss_ and lower D_ap_/D_loss_ (left and middle plots in **Figure 4D**). To confirm that this effect correlated with biopolymer composition, we quantified Pel and eDNA in those biofilms using WFL and TOTO-1. As shown by the right plot and fluorescence images in **Figure 4D**, the WFL/TOTO-1 fluorescence intensity ratio increased with increasing rhamnose concentration, indicating a higher Pel:eDNA ratio. Together, these *in vitro* and *in vivo* results demonstrate that the polymeric composition of the biofilm matrix modulates PYO retention and EET in a predictable manner.

**Figure S5.**
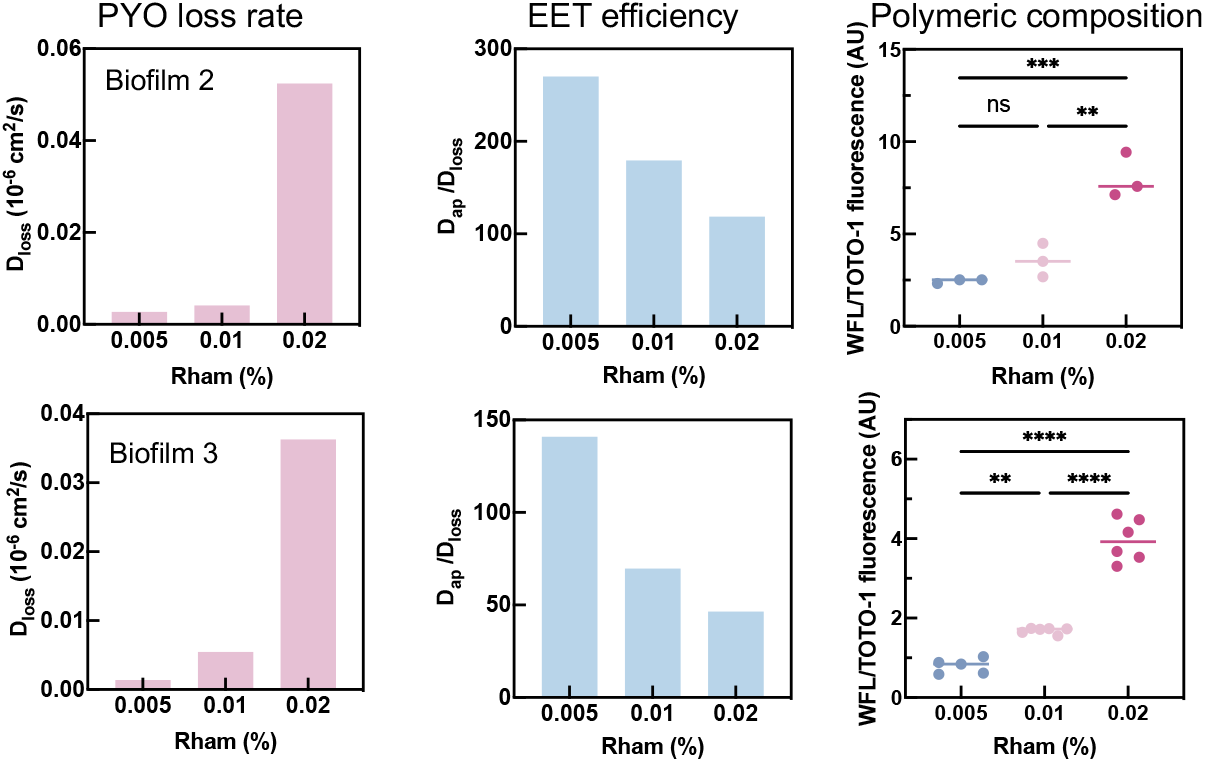
Triplicate datasets for Figure 4D, showing biofilm EET depends on Pel:eDNA ratio.

### Decreasing Pel abundance in the matrix enhances biofilm metabolism under O_2_ limitation

Next, we investigated how the matrix polymeric composition affects biofilm metabolism. *P. aeruginosa* uses EET to sustain the metabolism of O_2_-limited cells in the biofilm core by shuttling electrons from intracellular metabolism to distant electron acceptors (e.g., O_2_ at the biofilm periphery)^42–44^. Given that biofilms with a lower Pel:eDNA ratio retain more PYO and exhibit more efficient EET, we hypothesized that they would also display higher metabolic activity under standard conditions (i.e., oxygen-limited, illustrated in **Figure 5A**).

**Figure 5.**
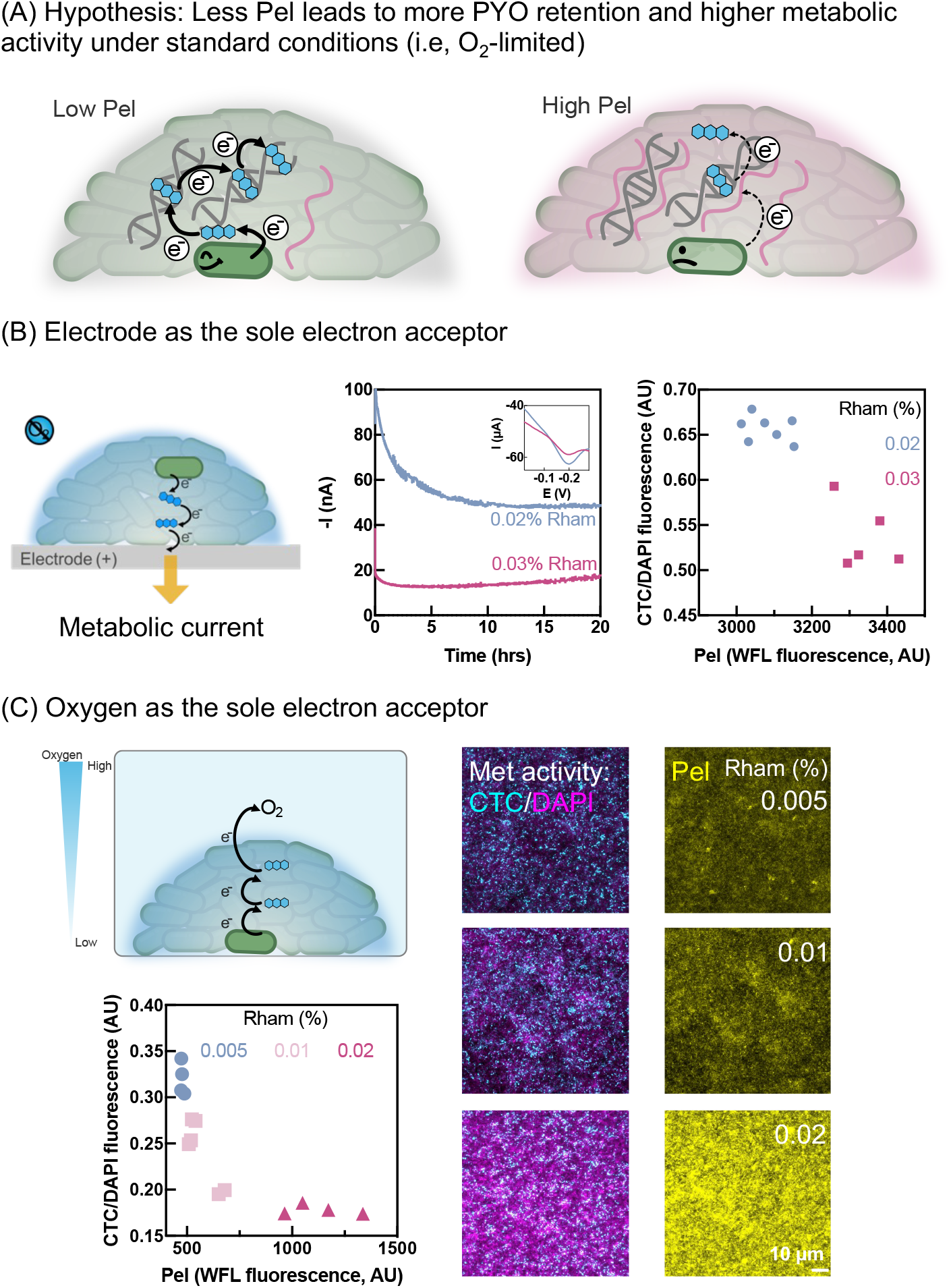
Less Pel enhances metabolism under O_2_ limitation. **(A)** Schematic: biofilms with less Pel retain more PYO, enabling more efficient EET and sustaining higher metabolic activity in O_2_-limited cells. **(B)** Electrode-based assay under anoxic conditions (schematic, left). Biofilms induced with lower rhamnose concentrations retained more PYO (inset in the middle) and generated higher metabolic currents (middle); analysis of confocal imaging shows biofilms induced with lower rhamnose concentrations are of less Pel and higher metabolic activities (right). Data are representative of biological triplicates, with triplicate datasets shown in Figure S6. At least three fields of view were imaged per sample; each dot in the right plot represents one field of view. **(C)** The oxygen-based assay confirms the same inverse relationship between Pel content and metabolic activity. Top-left: schematics of the experimental setup. Bottom-left: biofilms induced with less rhamnose exhibited higher metabolic activity (lower CTC/DAPI ratio) and reduced Pel. At least three fields of view were imaged per sample, and each dot represents one field of view. Data are representative of biological triplicates, with triplicate datasets shown in Figure S7. Representative fluorescent images from average intensity projections of Z-stacks are shown on the right.

We first tested this hypothesis using the electrode as the sole electron acceptor. As illustrated in **Figure 5B** and previously demonstrated^23^, an oxidative potential applied to the electrode drives electron transfer from intracellular metabolism via PYO-mediated EET, generating a measurable metabolic current. *P*_*rha*_*pel* biofilms were grown on gold chips in the anaerobic chamber under varying rhamnose concentrations to modulate Pel production. After thorough rinsing, biofilms were soaked in PYO, then transferred to fresh PYO-free medium (pH 6.5, anoxic) to mimic an open system where only biofilm-retained PYO persists. After 1 h, square-wave voltammetry confirmed that less-induced biofilms retained more PYO (inset of middle plot, Figure 5B). Subsequently, an oxidative potential (+0.1 V vs Ag/AgCl) was applied for 20 h; biofilms induced with 0.02% rhamnose generated a higher metabolic current than those induced with 0.03% rhamnose (middle plot, Figure 5B), consistent with our prediction.

To further validate the link between Pel content, PYO retention and metabolic activity, we characterized biofilms by confocal microscopy. Metabolic activity was assessed using 5-cyano-2,3-ditolyl tetrazolium chloride (CTC), a redox dye that is reduced intracellularly by metabolically active cells to insoluble fluorescent formazan.^45^ Pel was stained using WFL, as previously described. Biofilms were fixed with formaldehyde after staining and before removal from the anaerobic chamber for imaging. DAPI was used as a counterstain, and the ratio of CTC to DAPI fluorescence intensities served as a semi-quantitative measure of metabolic activity. As shown in the rightmost plot of **Figure 5B**, biofilms with lower Pel content exhibited higher metabolic activity, consistent with the metabolic current measurement.

Finally, to assess whether these results would extend to a more physiologically relevant condition where oxygen serves as the electron acceptor rather than an electrode, biofilms were grown in glass-bottom dishes in an anaerobic chamber, then exposed to oxygen to establish an O_2_ gradient across the biofilm.^46^ After thorough rinsing with PBS and soaking in PYO, the PYO solution was replaced with fresh medium to allow unretained PYO to diffuse out. Because PYO retention could not be directly quantified electrochemically in this setup, we instead measured PYO concentrations in the added and removed solutions by absorbance at 690 nm. The difference between these values indicated that for a representative biofilm induced with 0.005%, 0.01%, and 0.02% rhamnose retained 28%, 26%, and 23% of PYO, respectively. After 24 h of incubation, biofilms were stained with CTC, WFL, and DAPI as described, and the same inverse correlation between metabolic activity and Pel content was observed (**Figure 5C**).

Taken together, these results reveal that biofilms with less Pel exhibit higher metabolic activity under standard conditions when oxygen is limited, consistent with the model that polymeric composition regulates anaerobic metabolism through its effects on PYO retention and EET efficiency.

**Figure S6:**
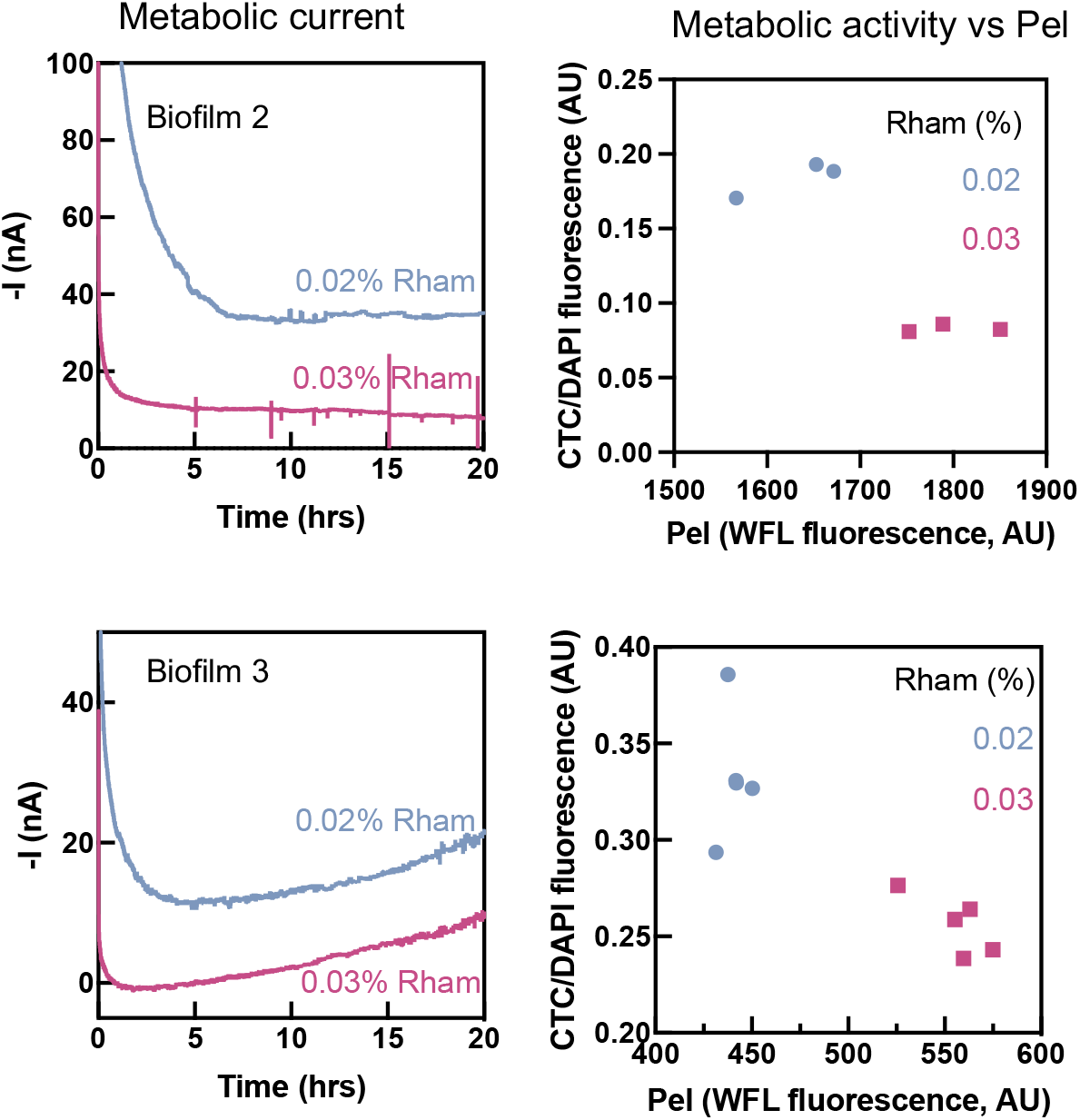
Triplicate datasets for Figure 5B, showing that less Pel enhances the metabolism of electrode biofilms.

**Figure S7:**
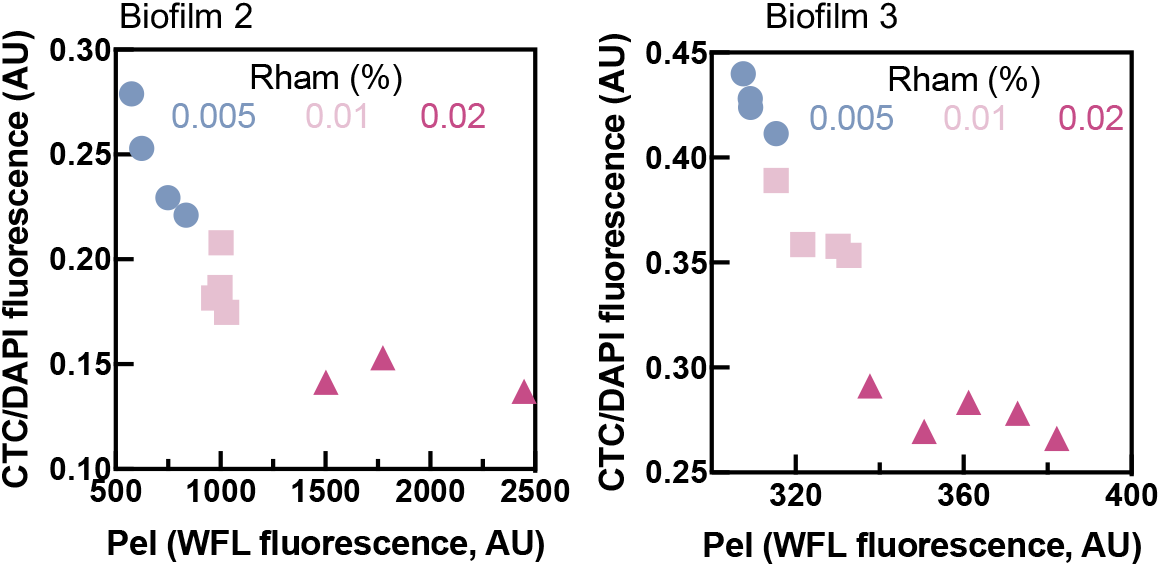
Triplicate datasets for Figure 5C, showing that less Pel enhances biofilm metabolism when oxygen is the sole electron acceptor.

### Increasing Pel abundance in the matrix protects biofilms from antibiotic treatment

Finally, having shown that Pel-rich biofilms exhibit lower metabolic activity, we wondered whether such polymer-driven attenuated metabolism might be beneficial in certain cases. Extensive research has linked biofilm antibiotic tolerance to low metabolic activity in the biofilm core, ^47–51^ as many antibiotics require active metabolism for uptake or action (e.g., targeting cell wall synthesis or protein production). We therefore speculated that modulating polymeric composition in a manner that slows metabolism would provide a survival advantage for cells challenged by certain antibiotics (**Figure 6A**).

**Figure 6:**
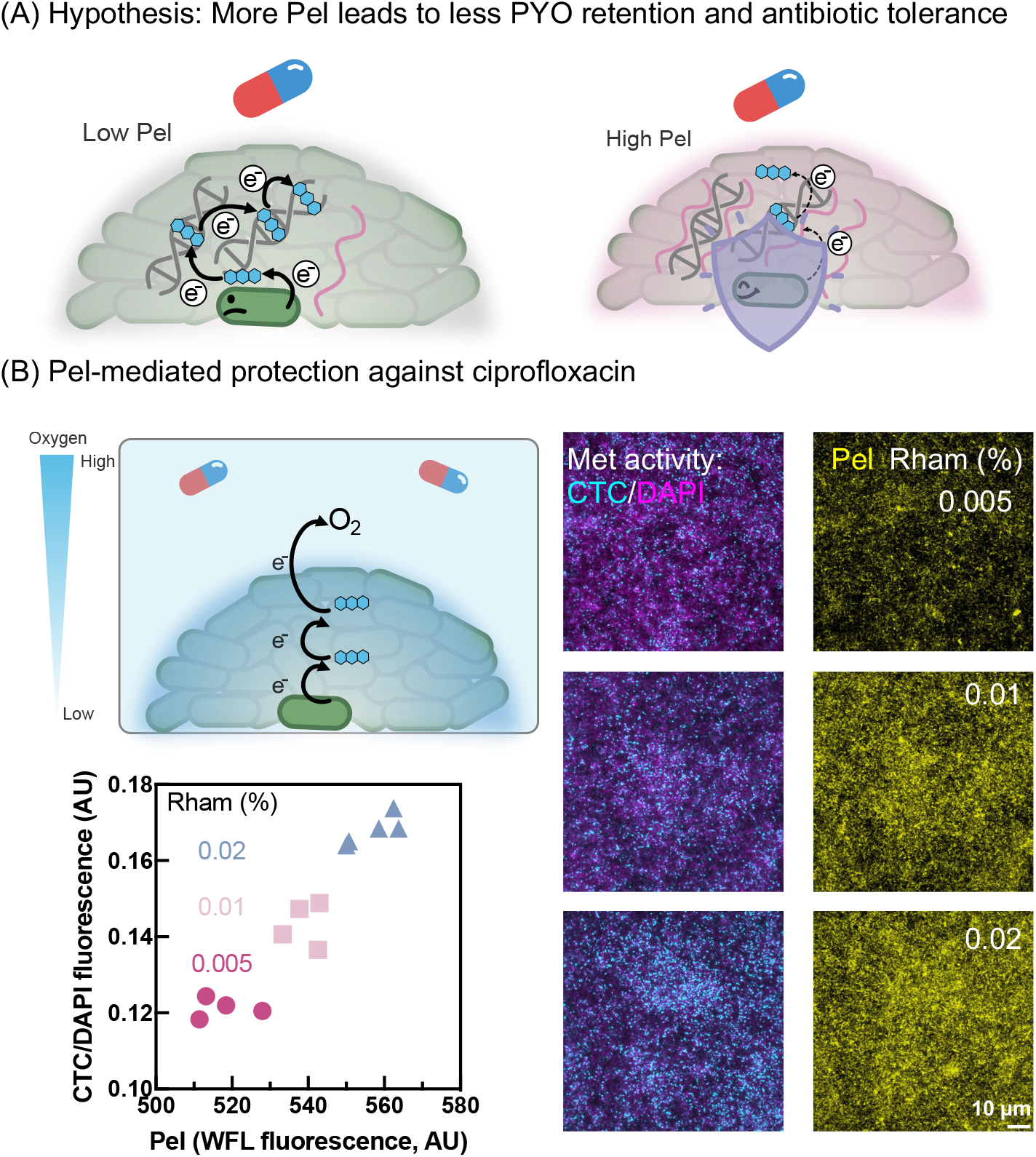
More Pel in the matrix protects biofilms from ciprofloxacin treatment. **(A)** Schematic: Pel-rich biofilms retain less PYO, resulting in increased ciprofloxacin tolerance. **(B)** Top-left: schematics of the experimental setup. Bottom-left: biofilms induced with more rhamnose showed enhanced survival (higher CTC/DAPI ratio) and more Pel. At least three fields of view were imaged per sample, and each dot represents one field of view. Data are representative of biological triplicates, with triplicate datasets shown in Figure S8. Representative fluorescent images from average intensity projections of Z-stacks are shown on the right.

To test this prediction, we stressed biofilms using ciprofloxacin, an antibiotic whose tolerance by *P. aeruginosa* biofilms is known to be linked to slow metabolism.^52,53^ Moreover, ciprofloxacin is uncharged at physiological pH^54^, minimizing electrostatic interactions with matrix polymers, enabling it to readily penetrate biofilms. This important chemical feature allowed us to focus on how tuning polymeric composition could impact antibiotic susceptibility via metabolic effects without influencing antibiotic penetration into the biofilm as previously described.^52,55^

Biofilms were grown in an anaerobic chamber. After removal from the chamber, they were rinsed with PBS, soaked in PYO, and then transferred to fresh medium containing 0.1 µg/mL ciprofloxacin but no PYO. After 4 h of incubation, biofilms were stained with CTC, WFL, and DAPI as described. As seen in the oxygen-limited biofilms without ciprofloxacin, increasing rhamnose inversely correlated with PYO retention (e.g. a representative biofilm induced with 0.005%, 0.01%, and 0.02% rhamnose retained 25%, 21%, and 18% of PYO, respectively). Biofilms induced with higher rhamnose concentrations exhibited higher Pel content and were more able to withstand the stress of ciprofloxacin, as reflected in their higher CTC/DAPI ratios compared to biofilms with lower amounts of Pel (**Figure 6B**). These findings suggest that while Pel enrichment imposes a metabolic disadvantage under oxygen-limited conditions (Figure 5C), it provides a metabolic advantage in the presence of ciprofloxacin, indicating that metabolic attenuation resulting from biofilm matrix effects have the potential to enhance antibiotic tolerance. Together, these results show that altering the polymeric composition of the biofilm matrix can influence cellular fitness.

**Figure S8:**
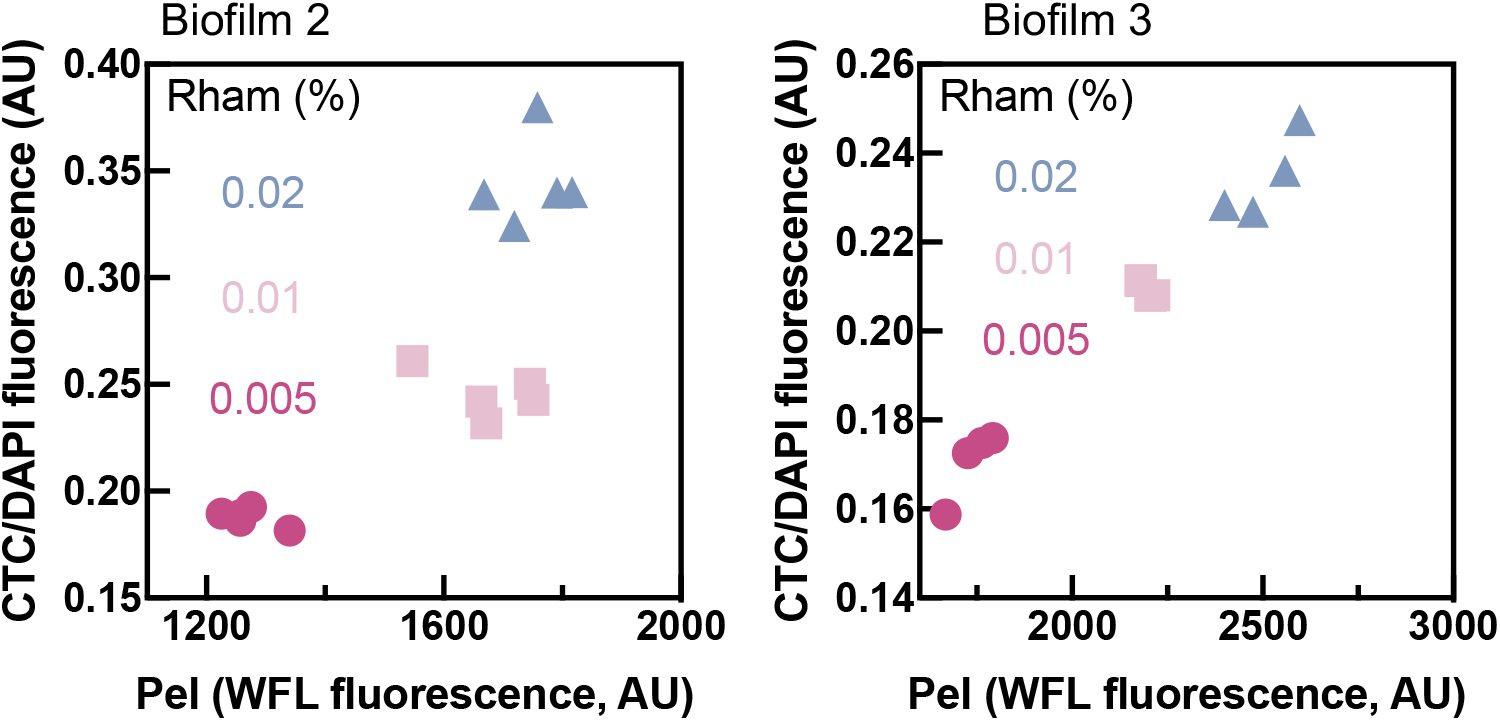
Triplicate datasets for Figure 6B, showing that more Pel protects the biofilm from ciprofloxacin treatment.

## Conclusion

While the effects of biofilm polymers on the mechanical properties of biofilms have been documented^37,56–58^, the metabolic effects of polymeric matrix chemistry that can be tuned as a result of biological activities have been largely unexplored. By applying materials science principles and engineering approaches to elucidate how matrix chemistry shapes biofilm physiology, we demonstrated that modulating the polymeric composition of the biofilm matrix can affect the biofilm metabolic rate, with important consequences for antibiotic tolerance. Specifically, we showed—using both biofilm-mimetic hydrogels and natural biofilms—that Pel and PYO electrostatically compete for eDNA binding in a pH and acetylation-dependent manner, which, together with the Pel:eDNA ratio, can have dramatic consequences for EET efficiency. Moreover, varying the Pel:eDNA ratio revealed a Goldilocks trade-off: lower ratios increased metabolic rate under oxygen limitation, whereas higher ratios enhanced survival under antibiotic stress.

These findings provide a proof of concept that polymer composition–property–function relationships in biofilms can be systematically studied and even predicted. This principle is likely generalizable well beyond the specific polymers and metabolites investigated here. Other strains of *P. aeruginosa* produce other exopolysaccharides, such as Psl (uncharged) and alginate (negatively charged). PA14 alone synthesizes four distinct phenazines with diverse chemical properties,^60^ and over 100 phenazine derivatives have been reported across other microorganisms.^61^ Could the same competitive and cooperative binding dynamics govern their EET efficiency? Could these principles explain why certain clinical strains dominate in specific niches?^62^ Beyond EET, we note that a variety of other processes, such as metal ion accumulation, could also be modulated by the composition of matrix polymers, as has been suggested for *Bacillus subtilis*.^63^ Our observation that distinct polymer ratios can provide cells with metabolic or survival advantages in different conditions suggests that cells may benefit from actively tuning the composition of their matrix by adjusting local pH, polymeric properties (e.g. extent of acetylation), or altering polymeric ratios. Elucidating whether and how such adjustments are achieved by different biofilm-forming species and how this may support different biological functions is an exciting goal for future research.

Beyond biofilms, our work offers a new conceptual framework for understanding the biological functions of extracellular matrices, in both prokaryotic and eukaryotic contexts. By treating the matrix as an integrated materials system, we highlight how the dynamic and complex interplay among matrix components shapes the local chemical environment that cells experience—governing the behavior of multicellular assemblies and driving emergent biological outcomes. For example, just as matrix composition controls PYO diffusivity, it may likewise regulate the transport and availability of other small molecules such as water, nutrients, metal ions, and other metabolites. Capturing the spatial and temporal dynamics of these interactions remains challenging but is essential for understanding the internal organization and function of multicellular systems.

### Experimental procedures

#### Strains and Strain Construction

Strains were derived from *P. aeruginosa* UCBPP-PA14 *Δphz\**, in which the *phzA1–G1, phzA2–G2, phzH, phzS*, and *phzM* genes are deleted.^23^ To construct Pel-inducible strain *P*_*rha*_*pel*, the native promoter region of the chromosomal Pel biosynthesis operon *pelA-G* was replaced with the rhamnose-inducible promoter *P*_*rha*_*BAD*. Markerless allelic exchange was performed using the plasmid pMQ30.^64^ Upstream and downstream DNA fragments flanking the native pel promoter were PCR-amplified from purified PA14 genomic DNA, while the rhamnose-inducible promoter was PCR-amplified from the pJM220 plasmid.^65^ The linearized pMQ30 backbone (SmaI-digested) and these fragments were assembled by Gibson assembly. The resulting plasmid was introduced into E. coli S17–1λpir and mobilized into the Δphz* strain by biparental conjugation. Following counter-selection on sucrose, the resulting *P*_*rha*_*pel* strain was verified by diagnostic PCR and sequencing. The *ΔpelA-G* mutant in a *Δphz\** background was engineered in a similar way, using a deletion plasmid provided by Dr. Lars Dietrich.^66^

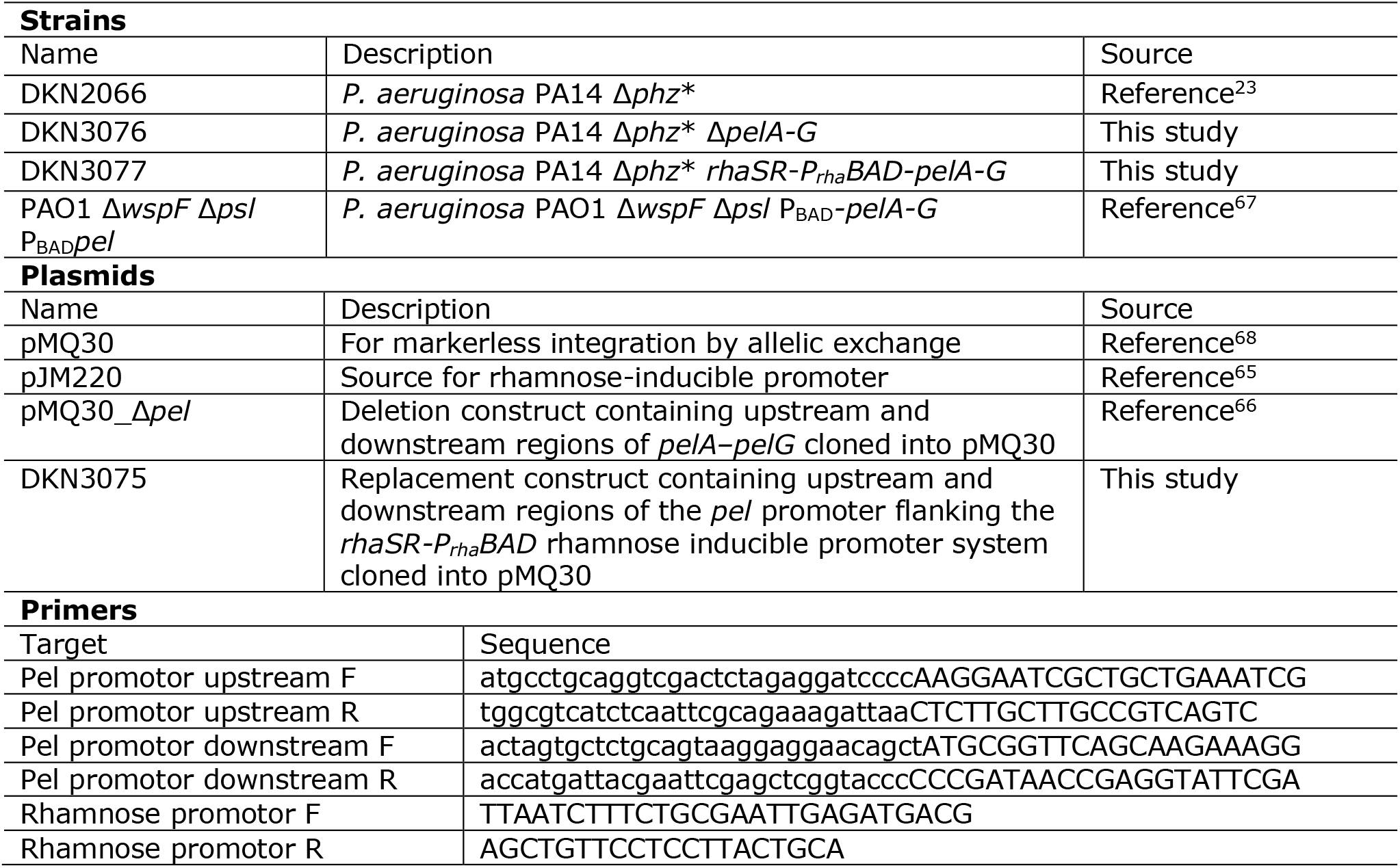

### Pel Isolation

As previously described^21^, Jensen’s medium supplemented with 0.5% arabinose was inoculated with 1 mL of overnight culture of the Pel overexpression strain PAO1 Δ*wspF* Δ*psl* P_BAD_*Pel*. Cultures were grown in baffled flasks for 20 h at 37°C with constant shaking. The supernatant was harvested by centrifugation (8,300 x g for 15 min at 20°C). Secreted Pel from culture supernatant was precipitated for 1 h at 4°C with ethanol (final concentration 75% v/v). The precipitate was washed three times with 95-100% (v/v) ethanol, resuspended in 20 mL of buffer (1 mM CaCl_2_ and 2 mM MgCl_2_ in 50 mM Tris, pH 7.5), and treated with 100 mg DNase I (Sigma Aldrich) overnight at 37°C, followed by 100 mg of proteinase K (NEB) overnight at 37°C. Sodium dodecyl sulfate (SDS) was added to the enzyme-treated samples at a final concentration of 4% (w/v) and heated to 95°C for 10 min. The Pel sample was centrifuged at 8,300 x g for 15 min at 20°C, and the resulting pellet was resuspended in water. The water washing step was repeated until the SDS was removed. The polysaccharides were precipitated again for 1 h at 4°C with ethanol (final concentration 75% v/v). The samples were extensively dialyzed against water (100 kDa molecular-weight cut off), and then flash-frozen and lyophilized for NMR analysis.

### Solid-state NMR spectroscopy

The solid-state NMR experiments were performed using a standard-bore Bruker Ascend magnet at 11.7 T (500.13 MHz for ^1^H and 125.78 MHz for ^13^C), a Bruker console with TopSpin software, and a 2.5-mm HXY magic angle spinning probe from Phoenix that was run in double resonance mode (tuning: H, ^1^H; X, ^13^C). Samples were spun at 11,111 Hz using a Phoenix PMAS unit. ^13^C CPMAS spectra were obtained with a contact time of 1.5 ms and a recycle delay of 3 s. Field strength for ^13^C cross-polarization was 50 kHz, and a linear ramp (ramp10090.100) was employed. ^1^H decoupling was at 100kHz (spinal64_p24) during acquisition. The ^13^C spectra were referenced to tetramethylsilane (TMS) as 0.0 ppm, which was determined relative to an external adamantane standard at 37.8 ppm. Line broadening of 80 Hz was applied. Multipeak fitting and integration were performed in software written for Igor Pro (WaveMetrics, Lake Oswego, OR, USA).

### Pel and PicoGreen Competition assay

Isolated Pel was dissolved in 20 mM HCl to a final concentration of 2 mg/mL by stirring overnight overnight at 4°C (final pH between 5 and 6). In black 96-well plates (Nunc, #237105), 100 µL of salmon sperm DNA (2 µg/mL; Sigma-Aldrich, #D1626) was mixed with 50 µL of PicoGreen solution (100-fold dilution from the stock; Thermo Fisher, # P11496) in 100 mM phosphate buffer. For the concentration-dependence assay, DNA and PicoGreen were mixed in pH 6.5 buffer, and 50 µL of Pel at varying concentrations was added to achieve the indicated final concentrations. For the pH-dependence assay, DNA and PicoGreen were first mixed in phosphate buffer at the indicated pH values, after which 50 µL of Pel was added to a final concentration of 20 µg/mL. In both assays, samples were incubated for 20 min in the dark before fluorescence was measured using a SpectraMax M2 plate reader. Following fluorescence measurement, the pH of each mixture was recorded to confirm that the introduction of Pel did not alter the buffer pH.

### NMR analysis of biofilm samples

^13^C CPMAS NMR spectra of the PA14 Δ*phz\** biofilm sample (containing biofilm matrix and bacterial cells) and the Δ*phz\** Δ*pel* bacterial cell sample were collected. The spectra were normalized to intensity at 21 ppm, which was chosen because it allowed for maximum subtraction of the Δ*phz\** Δ*pel* bacterial cell sample. The Δ*phz\** Δ*pel* bacterial cell spectrum was subtracted from the Δ*phz\** biofilm spectrum to obtain the difference spectrum, which was taken to be the spectrum corresponding to the matrix of the Δ*phz\** biofilm. This difference spectrum was subjected to multipeak fitting. The peak integrals for peaks located at 71, 84, and 101 ppm were used to estimate the relative glycan amount, and the peaks at 115, 129, and 138 ppm were used to estimate the relative DNA amount in the matrix of the Δ*phz\** biofilm.

Δ*phz\** biofilm was grown in an oxic bioreactor as previously described. ^23^ Briefly, a water-jacketed, three-ported glass reactor was used. The water jacket was connected to a heated recirculatory system (Polyscience, model 210) to maintain the temperature at 31ºC. The reactor was continuously stirred and aerated using an aquarium bubbler (Tetra #77851). One port served as the gas inlet, one as the gas outlet, and the third was used to suspend a glass slide in the reactor for biofilm harvesting. The reactor contained 100 mL of Jensen’s medium, inoculated to an OD_600_ of 0.1 from an overnight culture of Δ*phz\**. Medium was exchanged every 24 h. After 4 days, the biofilm-coated glass slide was removed, rinsed, lyophilized, and the biofilm was harvested for NMR analysis.

To prepare PA14 Δ*phz\** Δ*pel* cell samples for background subtraction, bacteria were plated on LB plates and grown overnight at 37ºC. A single colony was inoculated into 5 mL Jensen’s medium in borosilicate glass tubes and grown overnight at 37ºC with shaking. Overnight cultures were back-diluted to a final OD_600_ of 0.005 in 3 mL of Jensen’s medium in borosilicate glass tubes. Bacteria were grown statically at 25ºC for 72 hours. Bacterial cells were harvested via centrifugation, lyophilized, and analyzed by NMR.

### Preparation of Pel/DNA Hydrogels

A 2 mg/mL Pel stock solution was prepared as described above, and salmon sperm DNA (10 mg/mL) was dissolved in 200 mM phosphate buffer (pH 8.0). The two solutions were mixed to achieve the indicated Pel:DNA ratios, and appropriate volumes of each mixture were dispensed onto IDA electrodes, ensuring that the total DNA amount was constant across samples. After 1 h of drying in a 55 °C oven, the IDAs were soaked in pH 6.5 phosphate buffer for 1 min to allow gelation and rehydration. Electrochemical measurements were performed immediately afterward to quantify EET parameters.

### Growth of IDA *Δphz\** biofilms

*Δphz\** biofilms were grown in oxic bioreactors under conditions similar to those used for growing biofilms on glass slides for NMR analysis, with the exception that IDA electrodes were placed in the headspace of the reactor rather than immersed in the medium. This placement was chosen because aeration of the bioreactor produced fine droplets of medium in the headspace and provided higher oxygen availability compared to the liquid phase, creating favorable conditions for rapid biofilm formation. Biofilms were grown for 2–3 days before EET measurements were performed.

### Growth of *P*_*rha*_*pel* biofilms

*P*_*rha*_*pel* biofilms were cultivated in an anaerobic chamber on various substrates (e.g., IDA electrodes, gold chips, or glass) to obtain biofilms with different Pel:eDNA ratios. Overnight cultures of *P*_*rha*_*pel* were back-diluted to a final OD_600_ of 0.2 in LB supplemented with 100 mM KNO_3_ and 0.4% (w/v) rhamnose. For IDA electrodes and gold chips, 4 mL of culture was dispensed into each well of a 6-well plate containing a single IDA electrode or gold chip (Platypus Technology, #AU.1000.SLC_10mm). For glass substrates, 800 µL of culture was added to each well of a 4-well chambered coverglass (Nunc, #155382PK). Biofilms were first incubated anaerobically at 37 °C for 6 h to promote cell attachment, after which the medium was exchanged for LB containing 100 mM KNO_3_ and the desired rhamnose concentration. A second medium exchange was performed after 3 days. Following another 3 days of incubation, mature biofilms were obtained. Biofilms were rinsed thoroughly with PBS to remove residual KNO_3_ prior to further analysis.

### EET measurements

EET parameters were measured as previously published.^23^ Briefly, gel-coated IDA electrodes were soaked in 300 µM PYO solution, whereas biofilm-coated IDA electrodes were soaked in 100 µM PYO solution. All electrochemical experiments used IDA electrodes as the working electrode, an Ag/AgCl reference electrode (3 M KCl; Basi, #MF-2056), and a graphite rod counter electrode (Alfa Aesar, #14738). Square wave voltammetry (SWV) was performed until no further increase in current was observed, indicating complete PYO saturation. Electrodes were then transferred to a PYO-free, N_2_-purged reactor, and 14 consecutive cycles of generator–collector (GC) and SWV measurements were performed. All solutions were 100 mM phosphate buffer at the indicated pH. GC current was plotted against SWV current to generate a linear fit, with the slope used to calculate D_ap_. SWV current was plotted against time to calculate D_loss_.

### Metabolic current measurements

*P*_*rha*_*pel* biofilms were grown on gold chips with varying concentrations of rhamnose. Biofilm-coated gold chips were mounted on an L-shape platinum sheet electrode holder (Stony Lab, #SL000183) and immersed in 100 µM pyocyanin (PYO) at pH 6.5 until the SWV current no longer increased. The chips were then transferred to a PYO-free, N_2_-purged reactor containing 200 mL Jensen’s medium (pH 6.5, 70 mM glucose) to allow PYO to diffuse out. After 1 h, SWV was performed to assess PYO retention, followed by application of a constant +0.1 V potential for 20 h using the I–t technique to measure metabolic current. The reactor was subsequently moved into an anaerobic chamber, where CTC/DAPI staining was immediately conducted to evaluate metabolic activity.

### Metabolic activity measurements

Metabolic activity was quantified using the BacLight™ RedoxSensor™ CTC Vitality Kit (Thermo, #B34956) following the manufacturer’s instructions. Briefly, CTC working solution (15 mg/mL) was diluted 10-fold in PBS and added to biofilm samples. After incubation at 37 °C for 3 h in the dark, samples were rinsed and fixed overnight in 4% formaldehyde. Fixed samples were rinsed thoroughly and incubated with DAPI (25 µg/mL in water) for 30 min before fluorescence imaging.

### Polymeric composition characterization

WFL was fluorescently labeled with AZDye™ 647. WFL (5 mg; Vector Labs, #L-1350-5) was dissolved in 2.27 mL PBS (pH 7.4) to a final concentration of 2.2 mg/mL, and the pH was adjusted to 8 by adding 230 µL of 1 M sodium bicarbonate. AZDye™ 647 TFP ester (1 mg; Vector Labs, #FP-1129-1) was dissolved in 10 µL deionized water immediately before use and added to the WFL solution. The reaction mixture was incubated at room temperature on a rocker for 30 min in the dark. The reaction was quenched with 300 µL of 1 M Tris–HCl (pH 8.0) and incubated for an additional 15 min. Unreacted dye was removed using Amicon Ultra Centrifugal Filters (PBS as the exchange buffer) with three wash cycles. Protein concentration was determined using a Nanodrop spectrophotometer and adjusted as needed.

To determine the Pel:eDNA ratio, biofilms were incubated with fluorescent WFL (100 µg/mL) and TOTO-1 (1 µM) for 30 min, followed by thorough rinsing and immediate imaging. For simultaneous metabolic activity and composition measurements, biofilms were co-incubated with CTC and WFL for 3 h, fixed, and stained with DAPI.

### Microscopy and image analysis

Confocal microscopy was performed using an LSM800 microscope equipped with a 63× oil immersion objective. WFL and TOTO-1 were excited using 640 nm and 488 nm lasers, respectively; CTC and DAPI were excited using 488 nm and 405 nm lasers, respectively. Z-stack images spanning the full biofilm thickness were acquired and processed in FIJI. Average Z-projections were generated from all slices, and mean fluorescence intensity was quantified for each channel.

## Acknowledgements

This work was financially supported by NIH grant (2R01AI127850-06A1). GRS is a National Mah Jongg League Fellow of the Damon Runyon Cancer Research Foundation (DRG 2439-21). We thank Cameron E. Scantlin and Steven Woodhams for assistance with biofilm matrix composition characterization, Lars

Dietrich for providing the plasmid used to construct the Δ*pel* mutant, and members of the Newman laboratory for constructive discussions throughout the study. Imaging was conducted at the Caltech Biological Imaging Facility.

## Author contributions

J.L. and D.K.N designed the research; J.L. performed the experiments and analyzed the data; G.R.S contributed to the design and analysis of imaging experiments; K.D. and C.R. performed and analyzed NMR; M.R.P provided intellectual support; and J.L. and D.K.N wrote the manuscript, with editing from all authors.

## Supplemental information

